# Influence of environmental and biological factors on functional connectivity of the superior temporal sulcus in neonates scanned at term age

**DOI:** 10.1101/2024.05.07.592881

**Authors:** Charlotte Mancuso, Maxime Bacquet, Lucas Benjamin, François Leroy, Ghislaine Dehaene-Lambertz

## Abstract

The superior temporal sulcus (STS), one of the first sulci visible during brain development, is a key region for human communication, notably hosting linguistic functions in the left hemisphere. Fetuses and premature newborns already process external sound, but the auditory environment is vastly different *in-utero* and *ex-utero.* Does this have an impact on the development of the auditory and linguistic networks? To answer this question, we studied the functional connectivity of regions bordering the STS, delimited in each individual including full-term and preterm male and female neonates born at different gestational ages but all scanned at term. We found that in addition to the expected contralateral connectivity, various STS parts were more strongly connected to specific distant regions, revealing the typical auditory/linguistic division across the two banks of the STS reported in adults. Furthermore, the right posterior STS was more connected to the contralateral hemisphere than the left. Finally, sex and premature birth had effects on both STS volume and connectivity. Female neonates displayed a lesser left-right asymmetry in STS depth, and heightened connectivity from the left posterior STS compared to males. Most importantly, despite equivalent scan age, full-term newborns had deeper sulci compared with preterms, and local connectivity in the right temporal region increased linearly with gestation length. These results emphasize the impact of both sex and early auditory environment on the setting up of the cerebral networks that might contribute to explain the milder impact of premature birth on language in females.

## Introduction

The superior temporal sulcus (STS) possesses several remarkable features. It is one of the longest sulci in the human brain. It hosts the main systems of human communication, linguistic functions in the left hemisphere (Hickok & Poeppel, 2007) and voice recognition in the right hemisphere (Belin et al., 2000; Blasi et al., 2011). It is also one of the first sulcus to appear during gestation, with anatomical asymmetries immediately observed: The right STS appears earlier (Chi et al., 1977; Habas et al., 2012), is deeper than the left (Bartha-Doering et al., 2023; Glasel et al., 2011) as in almost all human adults (96% in Leroy et al’s report<u>, 2015)</u> and its width, particularly in the case of the right STS, has been related to enrichment with human gained enhancers (i.e. region of the genome which are active only in humans in interspecies comparisons of fetal brains) (Goltermann et al., 2024) confirming an evolutionary pressure on this structure.

Functionally, the auditory and linguistic networks in the superior temporal region are setting up during the last trimester of gestation, with brain responses to auditory input recorded since at least 30 wGA (Baldoli et al., 2015; Draganova et al., 2005; Jardri et al., 2008; Mahmoudzadeh et al., 2013; Moser et al., 2021). As the auditory environment to which fetuses and premature newborns are exposed is significantly different, we might wonder what impact it might have on the establishment of the auditory and linguistic networks. The *ex-utero* auditory world is enriched with higher frequencies, voices and speech are more distinct, but constant mechanical noise from medical equipment and the sudden shrill sounds of alarms might alter auditory development. In addition, frequencies in the voice range have been shown to be more attenuated in the incubator than in the womb (Monson et al., 2020) and duration of exposure to language is less important in preterm neonates compared to fetuses (2.6h/day vs 32mn/day reported by Monson et al., 2023).

These *in-* and *ex-utero* environmental differences might contribute to the higher risk of language deficits in children born prematurely than in those born at term, even in the absence of obvious brain damage (Guarini et al., 2009). Recent studies have indeed detected correlations between subtle anatomical disorganizations of the superior temporal region and its connections, and poorer language performance later in life: Aeby et al <u>(2013)</u> reported poorer language development in preterms whose MRI at term presented higher mean, transverse and longitudinal diffusivities in the superior temporal gyrus in the absence of any visible cerebral lesions. Salvan et al <u>(2017)</u> reported how at term age, microstructural properties of the arcuate fasciculus which links frontal and temporal linguistic regions were associated with later language development. Bartha-Doering et al <u>(2023)</u> observed that a smaller depth asymmetry between left and right STS in fetuses was associated with better verbal expression skills 10 years later and Kline et al <u>(2020)</u> reported a significant correlation between language scores at 2 years of age and the curvature of the left superior temporal gyrus (related to the STS depth) and insula, and surface of the left parietal in very preterm toddlers. Attempts to improve the auditory environment have been undertaken: Exposure to the mother’s voice produces an increase in the thickness of the auditory cortex measured by ultrasound (Webb et al., 2015) and an improvement in voice processing at term. Similarly, exposure to music improves musical processing at term (Adam-Darque et al., 2020; Lordier, Loukas, et al., 2019). In contrast, other studies have not observed changes in language acquisition milestones, such as responding specifically to the characteristics of the native language as a function of the duration of exposure to speech, but rather as a function of post-conceptual brain age (Pena et al., 2012; Peña et al., 2010). These latest studies emphasize the existence of biological constraints and critical windows within which exposure to speech must take place to allow the linguistic network to develop (Werker & Hensch, 2015).

In view of these studies, we sought to further explore STS development, notably its functional connectivity at full-term birth in order, on the one hand, to provide a better description of the early linguistic network and, on the other, to explore whether preterm birth might alter this connectivity. We also investigated the effect of biological sex, as a more asymmetrical brain, higher risk of cognitive impairment and slower development of communication skills tend to be reported in male individuals than in female ones (Etchell et al., 2018; Fenske et al., 2023; Hirnstein et al., 2019; Leroy et al., 2015; Williams et al., 2021a).

We capitalized on the data collected by the developing Human Connectome Project (dHCP; http://www.developingconnectome.org), which provides high-quality neonatal anatomical and functional MRI data. Because connectivity studies generally examine the voxel to voxel connectivity of the whole brain, they offer a global but locally coarse perspective, especially when it comes to a long structure such as the STS that encompasses various functional regions <u>(DeWitt & Rauschecker, 2012; Yeterian & Pandya, 1991; Dehaene-Lambertz et al., 2006; Pallier et al., 2011)</u>. Additionally, this sulcus exhibits complex twists and hemispheric asymmetries that pose normalization challenges for group-level analyses. Thus, to enhance precision, we meticulously reconstructed the shape of the STS in each individual and recovered the BOLD time-series in the contiguous cortical areas in each neonate. We subdivided each individual mask into four ROIs corresponding to anterior/posterior and inferior/superior divisions to create the seeds for functional connectivity analyses. After the correlations between the average BOLD oscillation within the seed and the whole brain were computed in each neonate’s native space and for each hemisphere, the connectivity maps were normalized toward a common template (Kabdebon et al., 2014) to allow comparison analyses. The division into superior and inferior portions was intended to distinguish the superior temporal gyrus, primarily involved in general auditory processing, from a lower region more implicated in linguistic functions (Dehaene-Lambertz et al., 2002; DeWitt & Rauschecker, 2012; Yeterian & Pandya, 1991), and so be able to reveal distant linguistic regions connected to the STS. Activations of the inferior frontal region have been observed in 3-month-old infants listening to short sentences (Dehaene-Lambertz et al., 2006) using fMRI, and in 30wGA preterm neonates discriminating syllables (Mahmoudzadeh et al., 2013). These observations suggest an early involvement of frontal regions in the human linguistic network, even in preverbal infants. On the other hand, other authors rather favor a progressive involvement with age of frontal areas (Perani et al., 2011; Skeide & Friederici, 2016). Looking at functional connectivity at birth in a large database should help to determine if the temporal receptive regions are already connected to frontal regions or whether this network is initially more segregated.

We also considered an anterior/posterior division to take into account the development of the STS depth asymmetry that might indicate a functional or ontogenetic parcellation associated with a specific connectivity (Bartha-Doering et al., 2023; Bonte et al., 2013; Leroy et al., 2015; Glasel et al., 2011). Indeed, the right STS is deeper than the left at the basis of Heschl’s gyrus. This asymmetry, called STAP for superior temporal asymmetrical pit, is observed in almost all humans (96% in Leroy et al’s report) but not in other primates, notably the chimpanzee (Leroy et al., 2015). A relation between voice activations and the deepest point of the STAP region has been shown (Bodin et al., 2018) as also a significant relation between the STAP anatomical barycenter and the functional barycenter for speech comprehension (Ardellier, 2018). The depth asymmetry increases in length along the STS towards the temporal pole from birth until adulthood (Falières, 2018; Leroy et al., 2015). We thus put the division point at the anterior limit of the neonatal STAP determined in a pilot study on dHCP release #0 (Falieres, 2018).

## Results

The regions of interest (ROIs) were defined for each neonate according to his/ her own anatomy, after delineating the shape of individual STS (figure 1A).

**Figure 1:**
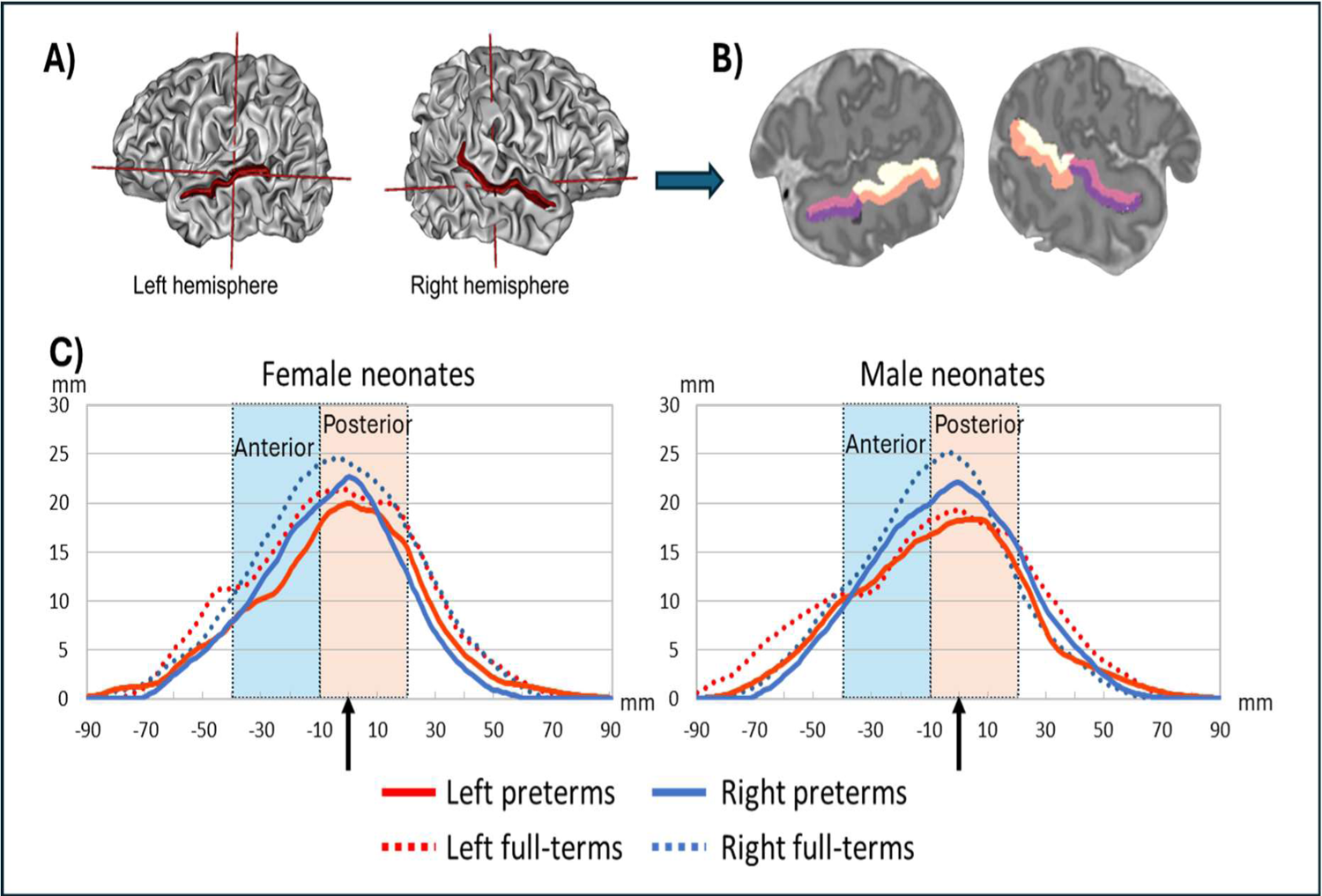
A) Identification of the STS in an individual neonate presented on her 3D mesh. B) its division into 4 ROIs presented on a sagittal view. The inferior and superior Rois correspond to the two banks of the STS and the anterior and posterior division was based on the boundary of the anterior neonatal STAP boundary (i.e. the region of asymmetric depth between the right and left STS, Leroy et al, 2015) determined on pilot data. C) Profile of the STS depth in male and female, preterm and full-term neonates. The coronal slice corresponding to the posterior tip of the planum temporale is chosen as the origin of the anterior-posterior scale (arrow). As expected, the right STS is deeper than the left and females are less asymmetrical than males. The STS is deeper in full-terms than preterm neonates in both hemispheres. The rectangles delimited the anterior and posterior limits of the ROIs.

### ROIs volumes

We first analyzed the ROIs volumes: there was a trend for larger ROIs in full-term than preterm neonates (F(1,112)=3.44, p=0.066), with a significant interaction group (full-terms vs preterms X antero-posterior ROI: F(1,112)=5.58, p=0.02). Furthermore, there was a significant interaction between sex and hemisphere (F(1,112)=7.37, p=0.008) and sex and antero-posterior ROI (F(1,112)=5.58, p=0.02). Figure 1C shows the average profile of STS depth in male and female neonates in preterms and full-terms. Sex and premature birth clearly affect the depth profile of the STS.

Post-hoc analyses showed that the right posterior ROI was significantly larger than the left, as expected due to the known deepest right sulcus (F(1,112)=8.31, p=0.005), the posterior ROI corresponding to the neonatal STAP (figure 1B-C). There was also an interaction sex X hemisphere for this posterior ROI (F(1,112)=9.27, p=0.003). The right-bias asymmetry was highly significant in male FT neonates (F(1,32)=17.02, p<0.001), but not in the other subgroups with the following order in the right-left volume difference: male FT > male PT > female FT > female PT. Significant pairwise comparisons (FDR corrected) were observed within male neonates between FT and PT (p=0.016) and within the full-terms between male FT and female FT (p=0.002). The increased asymmetry in full-term newborn males compared to females aligns with the same observation in adults (Leroy et al, 2015).

For the anterior ROI, there was an effect of group (F(1,112)=10.53, p=0.002) and of sex (F(1,112)=5.27, p=0.024) with no significant interaction between these two factors nor with hemisphere: Males had larger volumes than females, and full-terms than preterms. The difference between groups was due to the female preterm neonates who had the smallest anterior STS volume (female PT=504 mm3 < male PT=597 mm3 < female FT=643 mm3 < male FT=685 mm3). Significant pairwise comparisons (FDR corrected) were observed within the preterm neonates: female < male (p=0.020) and within female neonates: PT< FT (p=0.005). Interestingly, with regard to cranial perimeter, there was no main sex effect (F(1,112)=1.3, p=0.26), nor sex X group interaction (F(1,112)=1.29, p=0.26) at scan-time. A linear regression model aimed at predicting cranial perimeter from these two variables yielded non-significant results (F(2,113) <1). It was also the case for the left STS volume (F(2,113)= 1.88, p=0.16, r²= 1.5%) whereas both factors explained Right STS volume (F(2,113)= 10.69, p<0.001, r²= 14.4%, t=4.1, p<0.001 for term at birth and t=2.1, p=0.038 for sex). This suggests that sex and premature birth have a more specific effect on the right STS growth than on general brain growth.

### ROI-to-voxel analysis

For all analyses, significant clusters (p<0.001 at the peak level and p_FDR_<0.05 at the cluster level) are displayed in the corresponding figures. As expected given previous publications (Eyre et al., 2021), each ROI showed a significant correlation with the contralateral region (Figure 2). This correlation was not limited to the exact contralateral region but extended to the entire STS on the other side. Moreover, both left and right inferior ROIs displayed significant positive correlations with the BOLD signal in the inferior parietal and prefrontal areas and negative correlations with the region bordering the central sulcus. Right ROIs showed a strong correlation with the contralateral left regions while the left ROIs appeared mostly correlated with ipsi-lateral regions.

**Figure 2:**
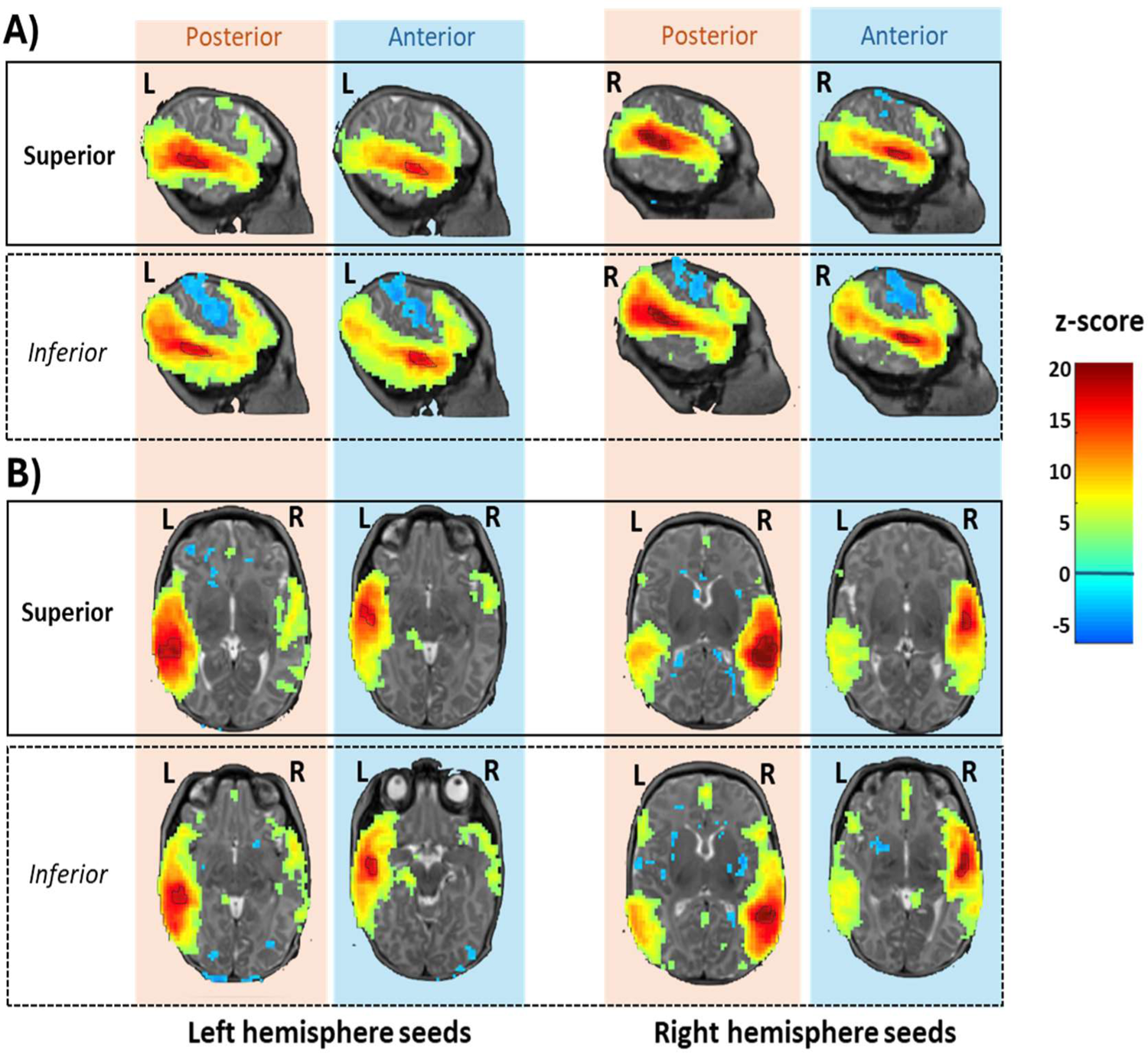
Functional connectivity of the left and right four ROIs. Clusters showing significant positive and negative correlations with the mean BOLD response of the ROIs (p<0.001 at the peak, p_FDR_ <0.05 at the cluster level) are displayed on sagittal (A) and horizontal slices (B) of an individual neonate. The chosen slices correspond to the maximum correlation peak in each analysis, trivially within the ROI analyzed. Significant clusters are seen in frontal areas, particularly for inferior ROIs, with significant negative correlations around the central sulcus. Note also on the horizontal slices (B) that contra-lateral correlation appears stronger for right ROIs (last two columns) than for left ROIs (first two columns).

### Comparisons between ROIs

#### Inferior vs Superior ROI (significant clusters are presented in figure 3A)

Trivially, each seed exhibited stronger correlations within itself (maxima of the blue and right clusters along the STS). Each ROI showed also stronger correlation with its specific contralateral region. But the most striking difference appeared in the more distant correlations: The superior temporal ROIs were significantly more correlated with the pre- and postcentral gyri bilaterally (blue clusters), whereas the BOLD signal from the inferior ROIs correlated with several regions in the inferior parietal and frontal lobes, along the dorsal peri-sylvian pathway (red clusters).

**Figure 3:**
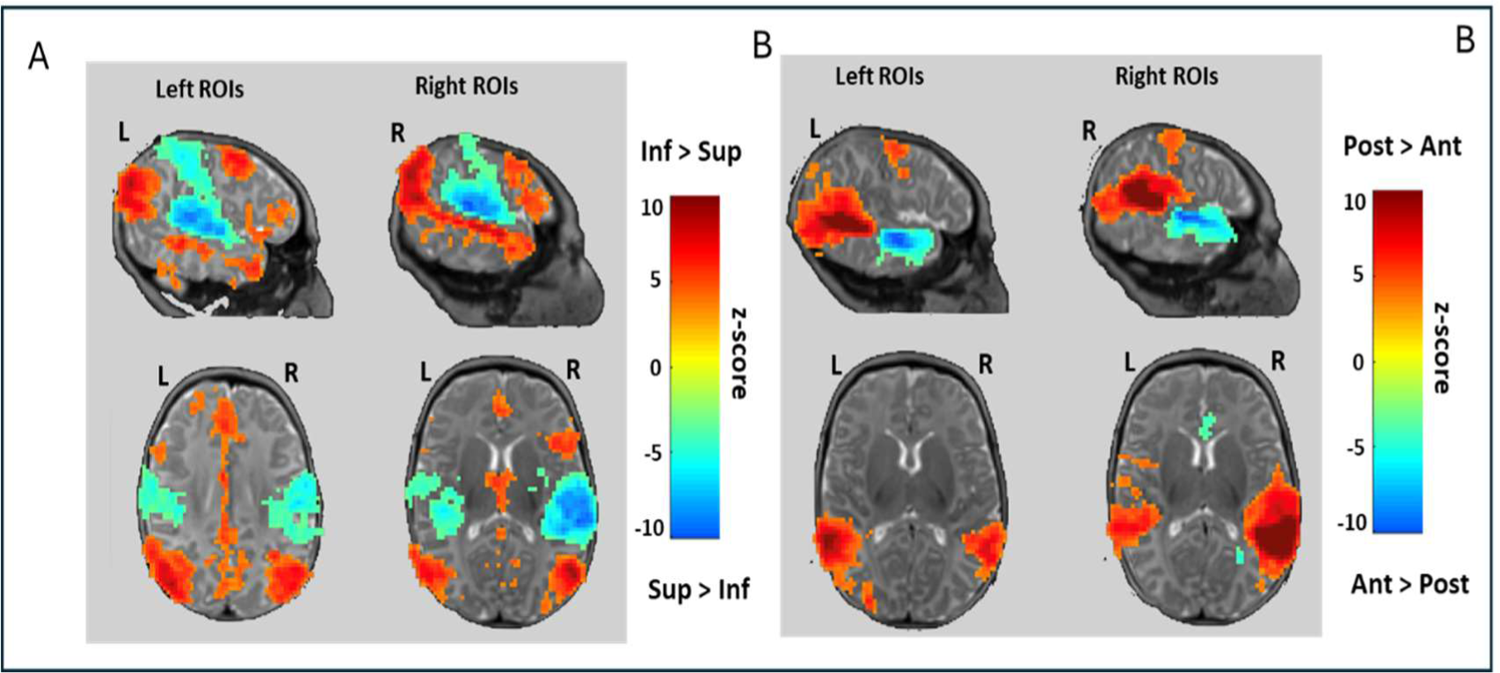
Differences in functional connectivity between superior and inferior ROIs (left) and between posterior and anterior ROIs (right) (p<0.001 at the peak, p_FDR_ <0.05 at the cluster level). The inferior ROIs are significantly more correlated with inferior parietal regions and prefrontal regions than the superior ROIs with a reverse effect in the central sulcus. Although the mean BOLD fluctuation in both anterior and posterior ROIs correlates with the contralateral STS as a whole, the posterior ROIs show a stronger correlation with their contralateral counterpart. The slices are selected at the coordinates of the peak difference.

#### Anterior vs Posterior ROIs (Figure 3B)

The posterior STS seeds were more correlated than anterior ROIs with the contralateral posterior regions regardless of whether the seed was on the left or right (Figure 3B, red clusters). By contrast anterior and posterior ROIs were similarly correlated with contralateral anterior regions. In other words, the mean BOLD response of the posterior ROIs was more in phase with the contralateral temporal region as a whole than was the case for the anterior ROIs. In addition, the posterior ROIs were more correlated with the precentral gyrus than the anterior ROIs (p_FDR_<0.001).

#### Left vs Right ROIs (Figure 4)

To overcome the anatomical asymmetries caused by the Yakovlevian torque and facilitate a comparison between the left and right hemispheres, we normalized the connectivity maps on a symmetrical space. Subsequently, we flipped the correlation maps issued from the right ROIs to compare them to the left correlation maps (see method). Compared to right ROIs, left ROIs extended their connectivity further back (27 voxels, p_FDR_ = 0.006, z=4.8 in the left angular gyrus; 34 voxels, p_FDR_ = 0.002, z=4.5 in the left posterior STS) and with the precuneus (47 voxels, p_FDR_ <0.001, z=4.48). Relative to left ROIs, right ROIs showed increased functional connectivity with the neighboring middle temporal gyrus (43 voxels, p_FDR_ =0.001, z=4.95) as with the contralateral posterior region (24 left voxels, p_FDR_ = 0.029, z=4.62).

**Figure 4:**
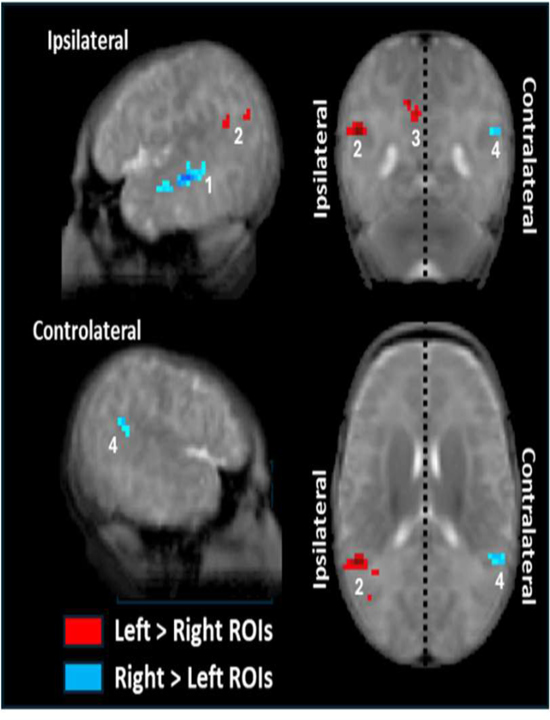
Differences in functional connectivity between left and right STS. Right and left ROIs were aligned on the same side of a brain that had been symmetrized to avoid alignment differences due to the Yakovlean torque. Thus, the left side of the brain corresponds to ipsi-lateral ROI connectivity and the right side to contralateral ROI connectivity, whereas the red color corresponds to clusters of stronger connectivity for left ROIs compared to right ROIs and the blue color to the opposite comparison. The right STS has stronger local correlation in its medial part (1) whereas it is with its posterior vicinity for the left STS (2) and precuneus (3). In addition, the right STS is more correlated than the left with the contralateral posterior region (4).

### Effect of sex and prematurity

After having characterized the functional connectivity of our ROIs, we performed linear analyses of their connectivity with sex and prematurity as regressors. In general, the significant effects were in the direction of stronger functional connectivity with longer gestation and in female than in male neonates.

#### Prematurity

We observed only positive correlations with gestation length. Considering the whole group, we observed a large linear effect of term for the right ROIs with stronger local connectivity in the right temporal region for infants born at a later term (Figure 5A) (anterior inferior: 200 voxels, p_FDR_ < 0.001, z=5.38; 27 voxels, p_FDR_ = 0.016, z=4.47; 37 voxels, p_FDR_ = 0.005, z=4.13; anterior superior: 621 voxels, p_FDR_ < 0.001, z=6.47; posterior inferior: 83 voxels, p_FDR_ < 0.001, z=4.96, 98 voxels, p_FDR_ < 0.001, z=4.78; posterior superior: 487 voxels, p_FDR_ < 0.001, z=5.13). This result remained in analyses restricted to each subgroup (full-terms and preterms). In addition, even within the full-term group, a longer gestation was correlated with an increase in connectivity of the posterior inferior ROI with the ipsi-lateral temporal pole (59 voxels, p_FDR_ < 0.001, z=5.31) and with the contralateral left posterior STS for both the inferior posterior ROI (39 voxels, p_FDR_ = 0.002, z=4.36) and the superior posterior ROI (72 voxels, p_FDR_ < 0.001, z=4.52).

**Figure 5:**
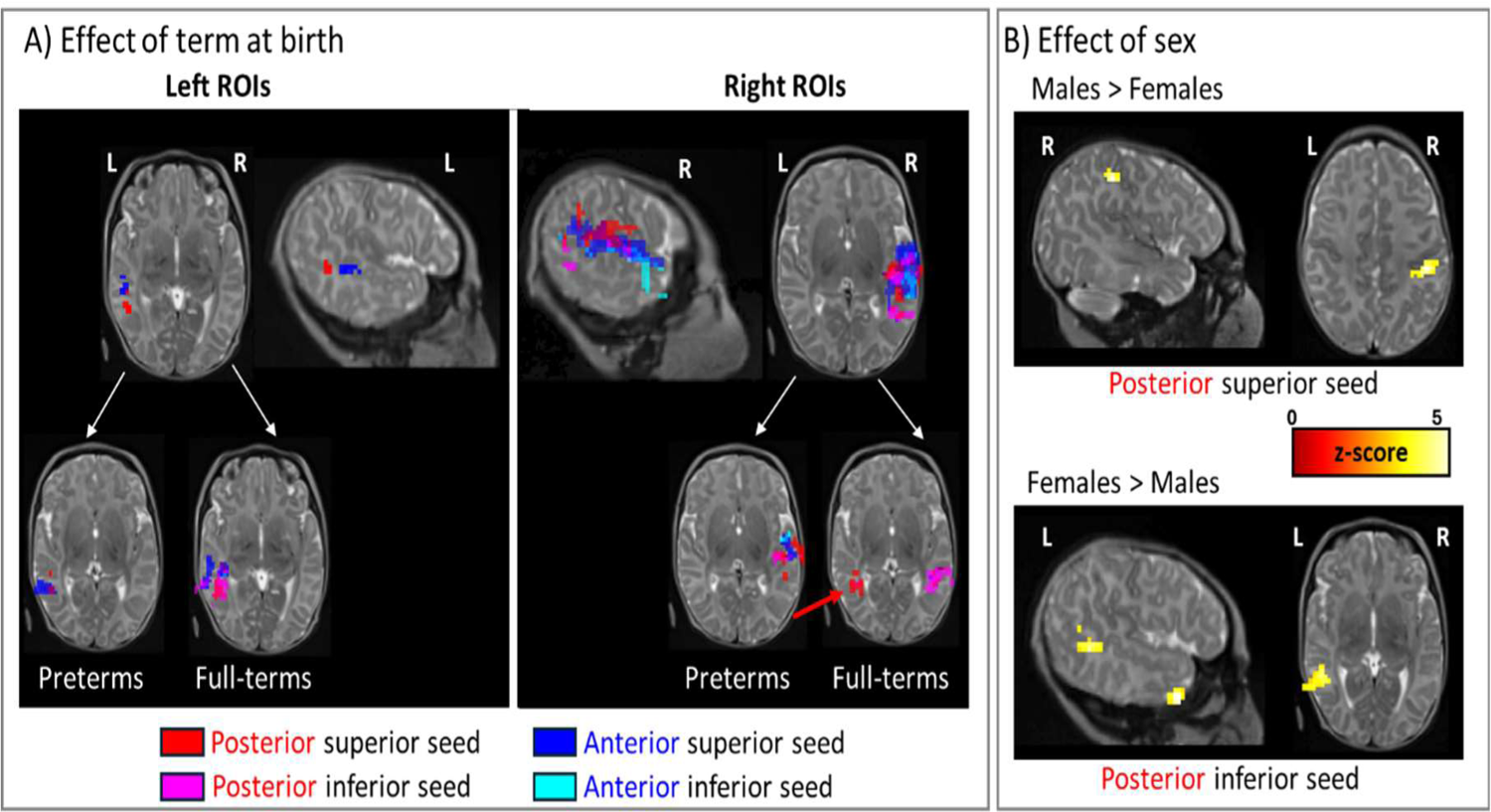
**A)** Positive effect of longer gestation on the functional connectivity of each ROI. Different colors correspond to the significant clusters in each analysis (p<0.001 at the peak level and p_fdr_ <0.05 at the cluster level), projected onto an individual newborn brain. Term at birth had a modest effect on functional connectivity for left ROIS contrasting with the sizeable effect for right ROIs whether the analysis concerned the whole group (top row) or each subgroup (bottom row). In addition, full-terms with a longer gestation display a stronger connectivity of the right posterior inferior ROI with the contralateral left region (red arrow). **B) Effect of sex on the functional connectivity of each ROI.** Male neonates showed stronger connectivity from the right posterior superior ROI to the ipsilateral postcentral gyrus (top row) whereas females showed a higher correlation between the left posterior inferior ROI with its inferior vicinity and the temporal pole (bottom row).

By contrast for the left ROIs, the effect of term at birth was modest, concerning only the superior ROIs (anterior: 26 voxels, p_FDR_ = 0.024, z=3.88 and posterior: 37 voxels, p_FDR_ = 0.005, z=3.82) when all neonates were considered. These results were also replicated in each subgroup (Figure 5A).

**Sex** (Figure 5B):

This factor mainly modulated the functional connectivity of the posterior ROIs. Notably female neonates displayed stronger local functional connectivity in the linguistic bank of the STS: from the left posterior inferior ROI to its above and below vicinity (98 voxels, p_FDR_ < 0.001, z=4.68 and 45 voxels, p_FDR_ = 0.001, z=4.47) and from the two left inferior ROIs to the temporal pole (36 voxels, p_FDR_ < 0.002, z=5.24 for the posterior ROI and 31 voxels, p_FDR_ = 0.013, z=4.66 for the anterior ROI). Male neonates had stronger correlations than females from the right posterior superior ROI to the postcentral region (32 voxels, p_FDR_ = 0.012, z=4.85).

The same pattern in favor of female neonates was observed in full-terms: the left inferior posterior ROI was more connected to its posterior vicinity (210 voxels, p_FDR_ < 0.001, z=4.85), and both posterior and anterior inferior ROI to the temporal pole (posterior ROI: 39 voxels, p_FDR_ = 0.004, z=4.63, anterior ROI: 38 voxels, p_FDR_ = 0.006, z=5.31). In addition, in the right hemisphere, the inferior anterior ROI was more connected to the temporal pole (34 voxels, p_FDR_ = 0.013, z=4.44) and the right inferior posterior ROI to its medial part (56 voxels, p_FDR_ = 0.001, z=5.22) and to the right hippocampus (30 voxels, p_FDR_ = 0.014, z=4.71) in females than in males. There was no significant cluster in the reverse direction. By contrast in preterms, there was no stronger functional connectivity for female relative to male neonates. The only difference being observed was in the reverse direction: the right superior posterior ROI was more connected to the right precentral region (41 voxels, p_FDR_ = 0.002, z=4.40) in male than in female preterm neonates.

## Discussion

The STS hosts important communication functions and undergoes important developmental changes during gestation, questioning how environmental factors might influence the setting-up of the auditory/linguistic network. To explore this, we investigated the connectivity of the STS using ROIs tailored to the specific STS morphology of each neonate. Comparing preterm to full term neonates, all scanned at term age, enabled to assess the impact of early acoustical environment on the development of this region. Figure 6 offers a visual synopsis of our key findings, which are listed below before we discuss their significance.

1. There was a striking difference in the functional connectivity between the two banks of the STS (figure 3A). The inferior bank was connected to inferior parietal and frontal areas, regions which, in the left hemisphere, belong to the classical linguistic network. This contrasts with the superior bank, whose BOLD modulations were more closely correlated with the areas bordering the central sulcus.
2. The posterior temporal region, notably in the left hemisphere, was connected with various nearby and distant regions, suggesting a functional hub role (Figures 3B & 4).
3. Premature birth had a notable impact on the growth of the STS, particularly in its anterior portion (FT > PT) despite measures being done at equivalent term age and confirmed observations in children (Zhang et al., 2015). As it can be seen in figure 1, the STS was deeper in full-terms compared to preterms. Additionally, the gestation length significantly influences the functional homogeneity within the right STS. Even within the full-term range, longer gestation increased the size of functionally homogeneous clusters in the right temporal lobe (Figure 4).
4. Male neonates compared to female neonates had a larger anterior STS and a more pronounced anatomical asymmetry at the level of the STAP region as it is also the case in adults (Leroy et al., 2015). Female neonates displayed a more extended connectivity of the left inferior ROIs (i.e. the linguistic STS bank) toward other temporal areas.

**Figure 6:**
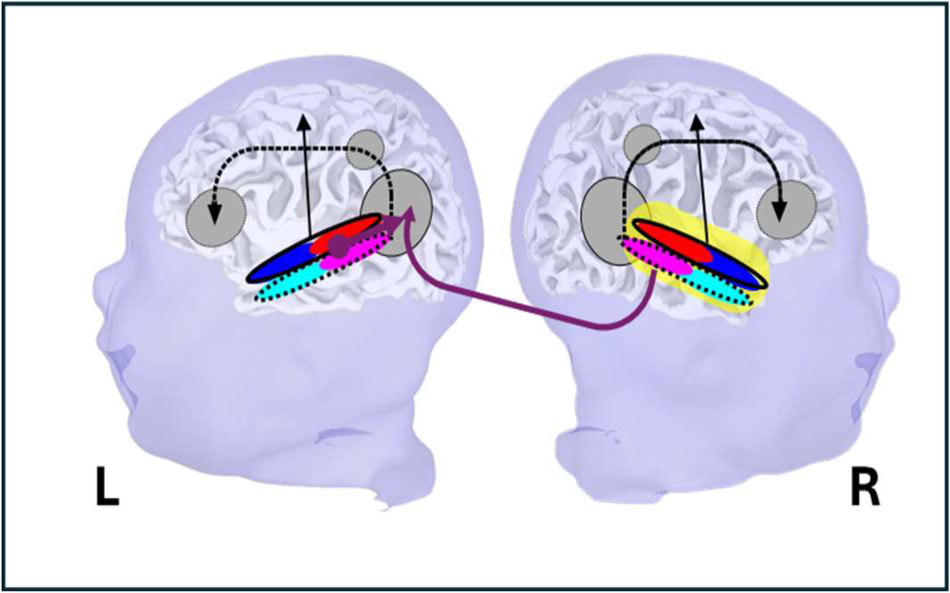
Schematic view of the functional connectivity of the STS in neonates. This functional architecture is roughly similar in both hemispheres except that the right STS is more connected to the left posterior region than the left to the right, a feature that is notably observed with longer gestation in the term range. The left posterior temporal region thus appears to be an important functional hub, whose connectivity is also affected by sex (stronger connectivity in female than male neonates). Eventually, the length of gestation has a significant and progressive effect on the local connectivity across the right superior temporal region (shown in yellow)

### The linguistic network at term age

#### Connectivity with frontal areas from birth-on

The clear connectivity of the inferior bank of the STS with regions along the dorsal linguistic pathway confirms the early setting-up of adult-like networks encompassing distant and high-level regions. It has been reported for other networks at the anatomical (<u>Vasung et al., 2019)</u> and functional levels using resting state studies (Barttfeld et al., 2018; Doria et al., 2010; Fransson et al., 2007, 2011; Kamps et al., 2020; Li et al., 2020; Sours et al., 2017; Sylvester et al., 2023). These results also converged with activation studies in post-term infants using fMRI (Dehaene-Lambertz et al., 2002) and before term at 29-30 wGA using NIRS (Mahmoudzadeh et al., 2013) that revealed frontal activations when infants are listening to speech stimuli. The increase in the number of participants as the increase in quality of the acquisitions due to progress in scanners and experience in infant data analyses provided by dHCP enable to obtain temporo-frontal correlations which were not easily seen in early studies <u>(such as Perani et al., 2011)</u>. This adds up to a growing body of evidence that frontal cortex is already functional and integrated in broader networks from birth (Mahmoudzadeh et al., 2013; Sylvester et al., 2023).

#### Left-right asymmetries in the posterior temporal region

As language is generally described as relying mostly on the left hemisphere, substantial hemispheric asymmetries might have been expected. To avoid spurious differences between hemispheres due to the Yakovlean torque that pushes up and forward the right hemisphere relative to the left, we have carefully created a symmetrical template to perform our left-right comparisons. Our results showed mostly comparable connectivity patterns in both hemispheres. This is in line with our own previous studies using speech stimuli in infants (Dehaene-Lambertz et al., 2002, 2006, 2010; Mahmoudzadeh et al., 2013), where we mainly observed bilateral activations, with even larger responses to speech syllables in most of the right hemisphere in preterm neonates, as it was also reported in Heschl’s gyrus in full-term neonates (Perani et al., 2011). Although hemispheric asymmetry in the language network undoubtedly increases with age (Emerson et al., 2016; Shultz et al., 2014), we have nevertheless consistently observed in our previous activation studies a left-hemispheric bias in the posterior temporal region. Here also, the left posterior temporal region appeared to occupy a peculiar position as a functional nexus connected to numerous regions.

It displayed stronger ipsilateral connectivity than the right, with its posterior and inferior vicinity, the superior part of the precentral region and the precuneus (Figure 3B-4). It is interesting to note that a female sex, as also a longer gestation, strengthened the connectivity of the temporal ROIs with the left posterior STS and middle temporal gyrus (figure 5). Finally, a longer gestation also accentuated the correlation of the right with this left posterior region suggesting an asymmetric inter-hemispheric transfer from the right to the left hemisphere (figure 5A bottom row).

An asymmetric inter-hemispheric between auditory regions through the corpus callopsum has been described in infants (Adibpour et al., 2020) and adults (Andoh et al., 2015; Gotts et al., 2013; Karpychev et al., 2022). In adults, repetitive stimulation of the right auditory cortex with TMS modulates its connectivity with the left contralateral temporal cortex but not the reverse. It has been explained by asymmetric callosal fibers favoring transfer toward the left temporal region (Andoh et al., 2015). Karpychev et al <u>(2022)</u> recently bring new argument to the role of the corpus callosum in language lateralization, showing the relation between language lateralization in adults and the volume of the temporal callosal fibers in the splenium measured with constrained spherical deconvolution (CSD). As the left posterior temporal region is notably involved in phonetic coding and intelligibility (Hickok & Poeppel, 2007; Vagharchakian et al., 2012), these results underline the centrality of this region in auditory-linguistic networks from the outset. The early and important role of the callosal inter-hemispheric transfer is also supported by the correlation between FA changes in splenium and linguistic performances in toddlers (Swanson et al., 2017). Children with greater acceleration in FA changes between 6 and 24 months in the posterior part of the corpus callosum, but not in classical language tracts, were greater talkers at 24 mo. Finally, the abnormalities often detected in the posterior corpus callosum in pre-term adolescents with language difficulties (Northam et al., 2012) might also be linked to the corpus callosum role to connect the right temporal regions to the left posterior temporal nexus.

The left posterior temporal region was also correlated with the precuneus in our analysis. The precuneus, a densely connected region is involved in many high-level tasks in adults, notably related to self-generated representations during rest. Therefore, its role might have been assumed to be negligible in infants. However, its functional connectivity already demonstrates in neonates a similar specificity than in adults (Sylvester et al., 2023) and it has been reported activated in 3month-old infants listening to their native language compared to backward speech (Dehaene-Lambertz et al., 2002). In adults, it is observed in the retrieval of items learnt during infancy, in association with the left temporal operculum (Fiebach et al., 2003), suggesting an early functional role of the precuneus that needs to be further explored during infanthood.

#### Auditory-sensory-motor connectivity

The sensory-motor regions bordering the central sulcus were observed in several of our connectivity comparisons and were mainly correlated with the posterior and superior ROIs (Figure 3 et 5B). This auditory-somatosensory connection, direct (Cappe et al., 2012) or mediated by the thalamus (Alcauter et al., 2014; Toulmin et al., 2015) is useful to plan the articulatory movement in function of the auditory return and might explain why a pacifier modulates auditory perception in infants (Choi et al., 2019, 2021). Thus even if the maturation of the dorsal pathway appears to be slightly delayed relative to the ventral pathway (Brauer et al., 2013; Dubois et al., 2009; Leroy et al., 2011), it is functional and might subserve phonological loops useful for short-term verbal memory (Dehaene-Lambertz et al., 2006) and early mapping between auditory, visual and motor speech representations (Bristow et al., 2008; Choi et al., 2019; Patterson & Werker, 2003).

Altogether, these results confirm the early setting-up of the linguistic pathways including frontal and sensory-motor regions and emphasize the role of the posterior temporal region and of the dorsal pathway long before infant’s production skills become apparent.

#### Effect of prematurity on the right STS functional connectivity

Beyond the strengthening connectivity with the left posterior temporal region reported above, the duration of gestation had mainly a linear impact on the functional connectivity of the right STS: Longer pregnancies correlated with larger clusters of homogeneous modulation of the BOLD time series in the right temporal lobe, an effect observed even in the restricted term range. This local connectivity is furthermore much greater in the right temporal than in the left across the entire group (figure 4). This result adds up to the list of observations showing faster development of the right hemisphere than the left: Most of the sulci, including the STS, appear earlier in the right than in the left hemisphere (Chi et al., 1977; de Vareilles et al., 2023; Habas et al., 2012); blood flow measured with single photon emission computed tomography (SPECT) is more important in the right hemisphere until 3 years of age (Chiron, 1997). Indices of maturation derived from T1, T2 MRI signal and DTI measures indicate more advanced development in in regions such as the right Heschl’s gyrus and *planum temporale* (Leroy et al., 2011). This fast development, notably during the age range considered here, might explain the stronger effect of environment on the connectivity of the right STS.

This environment effect on the right connectivity might be due to the specificities of the auditory environment, notably *in-utero*, where the abdominal barrier acts as a low-pass filter for external sounds (Querleu et al., 1988). Slow prosodic information is therefore the main characteristic of speech perceived in the womb, potentially favoring right auditory processes as right auditory regions have been described as more sensitive to slow frequencies (Boemio et al., 2005; Zatorre et al., 2004). This would explain the linear relationship between the right superior temporal connectivity and gestation length.

It is also congruent with the improvement of music and voice processing observed at term in preterms exposed to these stimuli compared to control groups (Adam-Darque et al., 2020; Lordier, Loukas, et al., 2019; Webb et al., 2015). Music and voice processing are biased toward the right hemisphere in infants and adults (Belin et al., 2000; Blasi et al., 2011; Bonte et al., 2013; Dehaene-Lambertz et al., 2010; Mahmoudzadeh et al., 2013) and these stimuli might be particularly efficient to maintain the development of the right STS functional connectivity in the *ex-utero* environment. Under this perspective, further studies are needed to better characterize the *in-utero* auditory world in order to design more targeted auditory interventions in the NICU, notably concerning language development.

#### Effect of sex on the posterior STS functional connectivity

The effect of sex on brain development and notably language is often discussed with mixed results (Christians et al., 2023; Etchell et al., 2018; Hirnstein et al., 2019). First, there are anatomical differences between male and female brain even when the sizes of the brain and the body are taken into account (Williams et al., 2021b). Second, girls have in average better language and social abilities than boys (Frank et al., 2017) and are less subjects to oral and written language pathologies (Altarelli et al., 2013). Finally, considering the effect of prematurity, it is classically estimated that girls present less negative consequences (Etchell et al., 2018; Hirnstein et al., 2019 but see the recent metanalysis of Christians et al., 2023) Although it is intrinsically difficult to disentangle cultural from biological factors, some studies report early biological differences in brain maturation and response to stress in males and females (Bale, 2016).

STS depth asymmetry was shallower in female than in male neonates, a feature already reported in adults. As males generally have larger brains than females due to their larger body size, the allometric relationship between gyrification and brain size (i.e. bigger brains are more gyrified) (Germanaud et al., 2012) might explain the sex difference in adults, but not here, given the lack of sex effect, as well as prematurity, on cranial perimeter. Despite this substantial anatomical difference, the effect of sex was moderate on functional connectivity. It consisted mainly of a stronger local connectivity from the posterior inferior ROI with other regions of the inferior bank of the STS (i.e. linguistic regions) in female neonates relative to males. Given the importance of these specific regions in language development, this more robust connectivity might play a role in girls better resistance to language pathologies.

The present study relied on a semi-automatic recognition of the STS morphology to be able to study specific networks. As the ROIs were based on individual STS with different length, depth and twists, the ROIs volumes were different (e.g. anterior vs posterior). This volume difference might have explained the differences in functional connectivity we report. However, the different results for inferior and superior ROIs rule out this explanation since superior and inferior ROIs sizes were similar by design. Thus, the observation of sex-related effects on the connectivity of inferior ROIs (but not superior ROIs) is not merely an artifact of ROI size but might reflect a stronger organization of the linguistic network in female neonates.

## Conclusion

Our study aimed at exploring the neural architecture of the STS at birth. By studying the impact of prematurity and sex, we hoped to discriminate as much as possible between the environmental and genetic contributions in the development of this region. The main divisions of the peri-sylvian (inferior vs superior, ventral vs dorsal and left vs right) networks are already observed at term age and remarkably poorly sensitive to a very different environment, except in the right temporal lobe. Those results highlight the need of studies in infants to investigate the neural bases of complex cognitive functions, such as language.

## Materials and methods

### Participants

All of the data used for the analysis were retrieved from a public database: “the developing Human Connectome Project dataset (dHCP) (release 2)” (www.developingconnectome.org). In this database, participants are preterm and full-term neonates, imaged at the Evelina Newborn Imaging Centre, St Thomas’ Hospital, London, UK. The document accompanying the release states that: “The study was approved by the UK Health Research Authority (Research Ethics Committee reference number: 14/LO/1169) and written parental consent was obtained in every case for imaging and data release”.

The release 2 of the database comprised 558 MRI sessions from 505 subjects, scanned between 24 and 45 wGA (see the release document for a complete description of the participants, http://www.developingconnectome.org/release-notes). We selected the infants scanned around term who got anatomical and functional runs with a good quality according to the quality control score given by a Senior Paediatric Radiologist published with the database. If there were twins, only one infant was randomly selected and we tried to span the entire premature range. We selected 159 subjects. After additional controls and difficulties in correctly extracting STS in some neonates, 116 subjects were finally retained.

After all data quality checking on the anatomical and functional data (see below), we obtained 116 participants (54.3% male neonates), comprising preterm (n=60) with a term at birth comprised between 24.3 and 37.9 wGA, and full-term neonates (n=56, term at birth >=38 wGA, max=41.7 wGA) (table 1). Among preterms, 31 were born before 35 wGA and 29 between 36 and 38 wGA. Preterms had no reported neurological impairment. All infants were scanned around term (mean=40.6 wGA [37-45.1]). At scan, term age and cranial perimeter were not significantly different across the two groups (table 1).

**Table 1:** Characteristics of the preterm and full-term groups (mean and range)

|  | **Preterms** | **Full-terms** | **P-value<sup>1</sup>** |
| --- | --- | --- | --- |
| n | 60 | 56 |  |
| gender (Male) | 30 (50%) | 33 (59%) | 0.3 |
| Term at birth (wGA) | 33.2 [24.3 - 37.9] | 39.7 [38.0 - 41.7] |  |
| birth_weight (kg) | 1.97 [0.54 - 4.10] | 3.24 [2.16 - 4.70] | <0.001 |
| Age (weeks) | 7.3 [0.1 - 17.1] | 1.0 [0.0 - 4.3] | <0.001 |
| Term at scan (wGA) | 40.42 [37.00 - 45.14] | 40.72 [38.14 - 44.86] | 0.4 |
| CP at scan (cm) | 34.40 [26.10 - 39.00] | 34.49 [30.50 - 38.00] | 0.8 |

<sup>1</sup> Welch two-samples t-test, except for sex analysis which consisted in a Pearson's Chi-squared test. Given the definition of the groups, the significant differences in birth weight and age were expected.

### Data acquisition

MRI was performed on a 3T Philips Achieva using a 32-channel phased array head-coil in a single session (63 mn) comprising structural and functional images during natural sleep (Hughes et al., 2017). We considered here the T2w anatomical image, because of its better signal to differentiate white and gray matter at that age relative to T1w, and the resting-state data images. The T2w images were acquired in axial and sagittal slices stack (0.8×0.8×1.6mm with a slice overlap of 0.8 mm; TR/TE=12000/156ms, SENSE factor=2.11). Functional images were acquired using multiband 9xaccelerated echo-planar imaging with high temporal resolution (TE/TR=38/392ms, voxel size=2.15×2.15×2.15 mm). The acquisition lasted for about 15 minutes (2300 volumes).

### Anatomical processing

T2w images were acquired and preprocessed by the dHCP team (Cordero-Grande et al., 2016). Brain mesh and sulci shapes were extracted and automatically labeled using the brainVisa software (Mangin et al., 2004; Perrot et al., 2011). M.B. visually inspected the results and corrected the labelling if necessary. Then, the deepest point of the *Planum Temporale* was located. The origin of the STS coordinates was determined as in Leroy et al (2015) and corresponded to the coronal plane showing the *planum* landmark. Eventually, the same parametrization as in Leroy *et al*. (2015) was applied to define a geodesic coordinate system along the STS including antero-posterior and inner-outer axes.

### Selection of the regions of interest (ROIs)

We restricted our analyses to the functional connectivity of the STS in the native space. Our seeds were thus created individually for each neonate from their own STS using the individual sulci shapes created as explained above (Figure 1). Then, they were dilated with a factor of 3mm in order to recover the cortical regions surrounding the sulcus, and finally binarized to create masks. We limited our analyses to the STAP region [-40, +20 mm] and delineated 4 ROIs in each hemisphere, corresponding to the superior and inferior banks of the STS and an anterior and posterior region of similar length along the STS to account for the progressive increase with age in length of the asymmetric segment towards the temporal pole. Indeed, the asymmetrical segment was 30mm-long in a preliminary analysis in 37 full-term neonates in the dHCP release 0 (Falières, 2018), 45mm-long in 3 months old and 55mm-long in adults (Leroy 2015). The asymmetry is visible in the posterior ROI [-10, +20mm] from term age onwards whereas it develops later in the anterior ROI [−40, −10mm].

The process of ROI construction was semi-automatic: all of the steps were double checked and potential mistakes due to the algorithm were manually corrected (e.g: label of another sulcus than the STS, inclusion of more than the horizontal part, inclusion of pieces of other sulci, or miscutting of the ROIs). Eventually, we were able to extract ROIs in 141 out of 158 neonates.

### Time series cleaning

Despite the motion correction performed during the preprocessing steps (Andersson et al., 2001, 2017, 2018), some data might remain noisy due to residual artefacts (magnetic field perturbation secondary to head movements and physiological processes, such as heartbeat, breathing *etc.*). To avoid sudden jumps that destroy correlation, especially between antero-posterior regions due to the typical pitch movement observed in infants, we applied the same method than Eyre *et al*. (Eyre et al., 2021), keeping only a continuous sub-sample of 1600 volumes, out of 2300 without movement ( ≈70% of the time series, ≈10 minutes of acquisition time) in each subject. We obtained 700 time-series. To choose the less noisy time series section, we used the movement parameters associated with the images (3 translation parameters and 3 rotation parameters). For each subject, we computed the framewise displacement (FD(t)) between each volume as the sum of the absolute values of the motion confounds. A threshold T was calculated for each time series, following the Eyre *et al*. method (Eyre et al., 2021), such as:

### IQR being the interquartile range, and C the value of the 75^th^ centile

Then, we simply calculated the number of points within the time series with a FD higher than *T*. This number corresponds to the number of outliers within the time series. Given that several time series had the same number of outliers, we ranked them on the sum of the distances between the threshold and FD(t) and retained the time series with the smallest combined distance between the threshold and FD(t). If more than 10% of the volumes (160 points) were outliers, the subject was excluded from the study. This method led to exclude 25 subjects. In the remaining 116 subjects, we finally applied a band-pass filter ([0.008-0.09] Hz) on the time series to remove the noise due to physiological processes (Yuen et al., 2019).

### Connectivity analysis at the subject level in the native space

All of the connectivity and statistical analyses were conducted in each infant’s native space using the python modules Nilearn (v.0.7.1) and Scipy (v.1.6.3). The time-series of all voxels in a ROI were averaged together to constitute the seed time-series. The seed-to-voxel correlation maps were computed across the whole brain using the Pearson coefficient (Mahadevan et al., 2020), and then z-scored. The mean time series of the white matter mask was also computed and included in the model as a non-interest confound.

### Normalization

All normalization steps were performed using SPM12 software (https://www.fil.ion.ucl.ac.uk/spm/software/spm12/). Once correlation maps had been calculated for each subject, all subjects were normalized to a reference space. We choose the infant template proposed by Kabdebon *et al*. (2014) because it has been parceled in anatomical regions following Tzourio-Mazoyer et al’s recommendations <u>(2002)</u>in the Automated Anatomical Labeling (AAL) atlas in adults and has been used in our other functional studies. To create a template reflecting our neonatal population, we first normalized toward Kabdebon et al’s template, 32 neonates, randomly selecting 8 neonates from each of 4 groups defined by their term at birth: <32, [32,35], [36,38], >=38. The obtained deformation field was applied to the grey matter, white matter, cerebro-spinal fluid and background masks distributed within dHCP (Draw-EM algorithm) and a mean image with these 4 compartments was obtained. All original T2w were then normalized to this mean image with a quadrilinear interpolation. The same deformation field was applied to the correlation maps to obtain all maps in the same reference space.

### Creation of a symmetrical template to study left-right functional asymmetries

In order to compare the connectivity from the left and right seeds, the correlation maps had to be registered in a perfectly symmetrical space, i.e. free of any hemispheric differences (petalia, Yakovlean torque, etc..). Thus, the infant template used for the previous group analysis was made symmetrical according to the procedure used by Didelot *et al*. (2010):

- The original template (orTemplate) was flipped along the x axis (fTemplate), and summed to the orTemplate. This creates a symmetrical image named orfTemplate.
- The orTemplate and fTemplate were coregistered on orfTemplate, then the mean of these two images was computed, creating the cTemplate.
- This last image was flipped again creating the fcTemplate. Finally, the mean of the cTemplate and fcTemplate was then computed to obtain the symmetrical template (sTemplate).

For each child, the following steps were then followed:

- The anatomical images were flipped along the x-axis
- The original and flipped images were normalized to the sTemplate created above and a deformation field was obtained for each image, which was applied on the corresponding left-seed correlation maps and flipped right-seed correlation maps. This assures that both images were aligned and had both been transformed through a normalization process toward the symmetrical template. The normalized left seeds correlation maps and flipped right seeds correlation maps were then compared.

### Statistical analysis

#### Anatomical ROIs

To analyze the anatomical features of the STS, we performed an Anova on the volumes of the masks obtained in each infant with 2 within-subject factors (anterior vs posterior location; left vs right hemisphere) and two between-subject factor of group (preterms vs full-terms) and sex. We further analyzed significant interactions by focusing on each ROI and group.

We also analyzed how term at birth (i.e. prematurity effect), civil age at scan (i.e. duration of exposure to the *ex-utero* life) and term at scan (i.e. brain “age”) might affect the volume of these ROIs in three different regression analyses. These regressors represent complementary ways of looking at the effect of brain maturation and environment, which are deeply intertwined. The effect of *ex-utero* environment should be better captured by civil age whereas term at scan should be more sensitive to brain biological maturation. Term at birth allows to see the effect of prematurity by itself.

### Functional analyses

All correlation maps were smoothed using a 3 × 3 × 3*mm* Gaussian kernel, then entered in second-level group ANOVAs using spm12. For the ANOVA-within subject analysis, the measurements were assumed to be dependent and the variance of each measurement considered as unequal. The same three factors than above were also considered in 3 different one-way ANOVAs for each seed.

Within-subject ANOVAs, with and without these regressors, were also performed to compare the connectivity between the different ROIs. Finally left-right comparisons were performed comparing the original connectivity images from the left ROIs and the flipped images from the right ROIs after both sides have been normalized to the symmetrical template to align both hemispheres despite the structural hemispheric asymmetries. Again, the same regressors were tested. Results are reported when voxels were significant at p < 0.001 and formed a contiguous cluster whose extent was significant at p<0.05, FDR corrected for multiple comparisons across the entire brain volume.

## Acknowledgments

The authors have received funding from the European Research Council (ERC) under the European Union’s Horizon 2020 research and innovation program (grant agreement No. 695710). We thank Mathilde Falières for her work on DHCP0 data during her master 2 internship.

## Declarations

### Availability of data and material

Data were provided by the developing Human Connectome Project, KCL-Imperial-Oxford Consortium funded by the European Research Council under the European Union Seventh Framework Programme (FP/2007-2013) / ERC Grant Agreement no. [319456] and ERC Grant Agreement no. [695710] to GDL and ENS de Lyon to CM. We are grateful to the families who generously supported this project.

### Code availability

Analyses codes are available upon request.

### Author’s contribution

GDL conceived the project. MB checked the quality of the data for inclusion in the study. LB, MB and FL performed the structural analyses. CM performed the functional analyses and CM, LB, FL & GDL wrote the paper.

